# Structural centrosome aberrations promote non-cell-autonomous invasiveness

**DOI:** 10.1101/216804

**Authors:** Olivier Ganier, Dominik Schnerch, Philipp Oertle, Roderick Y. H. Lim, Marija Plodinec, Erich A. Nigg

## Abstract

Centrosomes are the main microtubules organizing centers of animal cells. Although centrosome aberrations are common in tumors, their consequences remain subject to debate. Here, we studied the impact of structural centrosome aberrations, induced by deregulated expression of Ninein-like protein (NLP), on epithelial spheres grown in Matrigel matrices. We demonstrate that NLP-induced structural centrosome aberrations trigger the escape (’budding’) of living cells from epithelia. Remarkably, all cells disseminating into the matrix were undergoing mitosis. This invasive behavior reflects a novel mechanism that depends on the acquisition of two distinct properties. First, NLP-induced centrosome aberrations trigger a re-organization of the cytoskeleton, which stabilizes microtubules and weakens E-cadherin junctions during mitosis. Second, atomic force microscopy reveals that cells harboring these centrosome aberrations display increased stiffness. As a consequence, mitotic cells are pushed out of mosaic epithelia, particularly if they lack centrosome aberrations. We conclude that centrosome aberrations can trigger cell dissemination through a novel, non-cell autonomous mechanism, raising the prospect that centrosome aberrations contribute to the dissemination of metastatic cells harboring normal centrosomes.

## Introduction

Centrosomes function in the organization of microtubules and in ciliogenesis (Bornens, 2012; Conduit et al., 2015; Prosser & Pelletier, 2017; Sanchez & Dynlacht, 2016; Nigg & Holland 2017), and dysfunctions of these organelles have been linked to several human diseases, notably ciliopathies and microcephaly or dwarfism (Bettencourt-Dias et al., 2011; Braun & Hildebrandt, 2017; Nigg & Raff, 2009). Centrosome aberrations are also prominent in cancers, including preinvasive *in situ* carcinomas (Guo et al., 2007; Lingle et al., 2002; Pihan et al., 2003), suggesting that they contribute actively to carcinogenesis (Godinho & Pellman, 2014; Gonczy, 2015; Lingle et al., 1998; Nigg, 2002; Pujana et al., 2007; Zyss & Gergely, 2009). Best documented is the influence of centrosome aberrations on chromosomal instability, a hallmark of cancer (Ganem et al., 2009; Silkworth et al., 2009), and increasing evidence also suggests an impact on tissue architecture (Godinho & Pellman, 2014; Kazazian et al., 2017; Nigg, 2002; Raff & Basto, 2017). However, centrosome aberrations are generally present in only a fraction of all tumor cells and, moreover, expected to impair cell viability. Hence the question persists of whether and how centrosome aberrations contribute to cancer in humans (Chan, 2011).

Centrosome abnormalities are subdivided into numerical and structural aberrations, but in tumors the two defects are similarly prominent and often occur together (Guo et al., 2007; Kronenwett et al., 2005; Lingle et al., 1998; Lingle & Salisbury, 1999). The consequences of numerical aberrations have been studied extensively, culminating in the demonstration that centrosome amplification is sufficient to trigger tumorigenesis in both flies (Basto et al., 2008) and mice (Levine et al., 2017; Sercin et al., 2016). In contrast, structural centrosome aberrations have received little attention. They are presumed to reflect deregulated gene expression (Guo et al., 2007; Lingle et al., 1998), as illustrated best by NLP (Ninein-like protein), a centrosomal component implicated in the nucleation and anchoring of microtubules (Casenghi et al., 2003). NLP is commonly overexpressed in human tumors, notably in breast cancers and lung carcinomas, and overexpression of NLP promotes tumorigenesis in mice (Qu et al., 2008; Shao et al., 2010; Yu et al., 2009). When overexpressed to comparable levels in 2- (2D) or 3-dimensional (3D) cultures, NLP triggers structural centrosome aberrations that closely resemble those seen in tumor sections (Casenghi et al., 2003; Kronenwett et al., 2005; Salisbury et al., 1999; Schnerch & Nigg, 2016).

Recently we have shown that prolonged overexpression of NLP disturbs epithelial polarity and severely disrupts the architecture of spheres (Schnerch & Nigg, 2016). Here, we have used similar 3D models to explore the short-term impact of structural centrosome aberrations, with particular focus on the ability of epithelial cells to develop invasive behavior. We found that epithelia harboring NLP-induced centrosome aberrations trigger the selective dissemination of mitotic cells into the surrounding matrix and demonstrate that this novel type of invasive behavior is triggered by a non-cell-autonomous mechanism involving two concomitant processes: first, an impairment of E-Cadherin junctions and, second, an increase in cellular stiffness, which introduces heterogeneity into the biomechanical properties of the epithelium. These observations demonstrate that centrosome aberrations can trigger the dissemination of dividing cells from epithelia, and they suggest a novel mechanism for the inception of a centrosome-related metastatic process through multicellular cooperativity.

## RESULTS

### Structural centrosome aberrations trigger budding of living cells from epithelia

To determine whether centrosome aberrations influence the ability of cells to escape from an intact epithelium, we examined 3D acini grown in Matrigel, a natural hydrogel that mimics a basement membrane-like matrix (Artym, 2016). Remarkably, MCF10A-derived acini harboring NLP-induced structural centrosome aberrations frequently showed dissemination of individual cells into the surrounding matrix, a phenotype hereafter termed ‘budding’ (Fig 1A). Cell budding was observed in nearly 30% of MCF10A acini expressing NLP, whereas it was only seen sporadically (< 5%) in absence of transgene induction, or in acini expressing the control protein CEP68 (Fig 1B). Also, no budding was seen in response to overexpression of PLK4, suggesting that this phenotype is triggered primarily by structural rather than numerical centrosome aberrations. The same phenotype was seen with cysts prepared from Madine Darby canine kidney (MDCK) cells, attesting to its robustness (Fig 1A, Fig EV1). As judged from staining for cleaved-caspase 3, a marker for apoptosis, more than 65% of cells budding from NLP-expressing acini were alive, while more than 80% of the rare cells budding from control acini were undergoing apoptosis (Fig 1C-D). Collectively, these results show that structural centrosome aberrations induce the budding of living cells from the basal surface of polarized epithelia toward the surrounding matrix, potentially triggering an invasive process.

**Figure 1:**
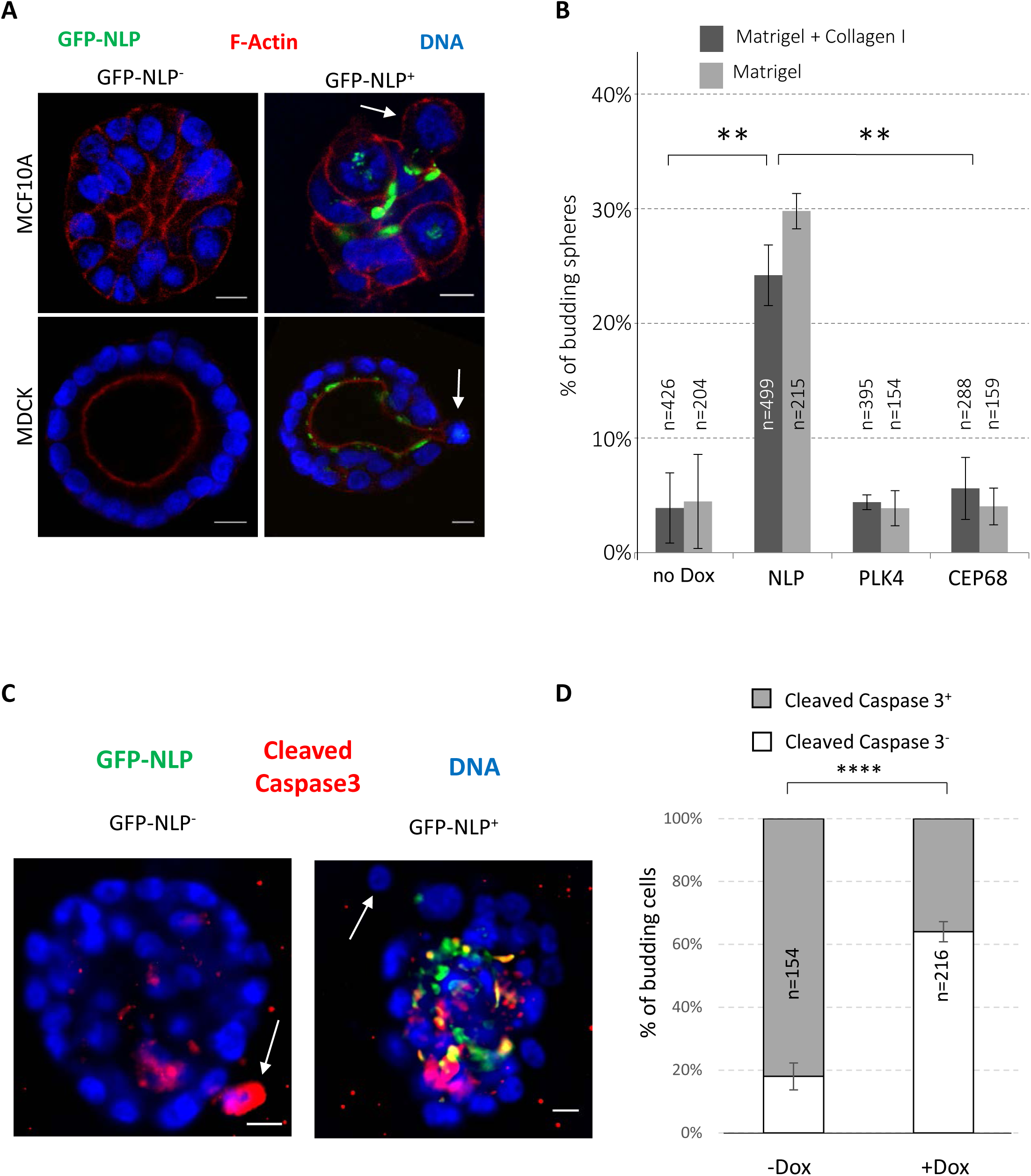
Structural centrosomal aberrations cause single cell budding from 3D acini. A. Representative images show acini derived from MCF10A (upper panels) or MDCK cells (lower panels), with or without induction of GFP-NLP expression. GFP-NLP^+^ and GFP-NLP^−^ acini were stained for F-actin (red) and DNA (blue). Arrows point to budding cells. Scale bars = 10 µm. B. Fraction of MCF10A acini that show budding in response to the indicated transgene products (GFP-NLP, GFP-PLK4 or GFP-CEP68), depending on whether acini were cultured in collagen I-enriched Matrigel (dark grey) or pure Matrigel (light grey); n indicates sample size and error bars indicate ± standard deviation (s.d.) of the mean from 3 independent experiments. (**) indicates a P-value <0.01, as derived from unpaired, two-tailed Student’s t-test. C. To determine the proportion of apoptotic cells, MCF10A acini expressing GFP-NLP (+Dox) or not (-Dox) were fixed and stained for cleaved caspase 3. Scale bars = 10 µm. D. Fraction of cells budding from MCF10A acini that are positive (grey bars) or negative (white bars) for cleaved-caspase 3 staining; data are shown for acini with (+Dox) or without (-Dox) induction of GFP-NLP expression. n indicates number of budding cells analyzed; error bars indicate ±s.d. of the mean from 3 independent experiments. (****) indicates a P-value <0.0001, as derived from unpaired, two-tailed Student’s t-test.

We emphasize that the budding phenotype is distinct from the formation of invasive protrusions (invadopodia) that occurs in MCF10A acini in response to PLK4-induced centrosome amplification (Godinho et al., 2014). In agreement with Godinho and coworkers, we also observed invadopodia formation upon overexpression of PLK4 (Fig EV2), provided that the invasion assay was sensitized by addition of type I collagen to Matrigel (Artym, 2016; Godinho et al., 2014; Schnerch & Nigg, 2016). However, we also observed invadopodia formation with acini harboring NLP-induced structural centrosome aberrations (Fig EV2), suggesting that this phenotype may represent a more general response to centrosome aberrations than hitherto assumed.

### Budding involves mitotic cells

Immunofluorescence staining suggested that most, if not all, budding cells were going through mitosis (Fig 2A). To corroborate this conclusion, we performed time-lapse microscopy on MCF10A acini and MDCK cysts derived from cells that stably expressed mCherry-α-tubulin. NLP overexpression was induced in acini and cysts after 5 days, and these were then imaged every 20 minutes for 3 days, resulting in the recording of 35 independent budding events. Remarkably, all these budding events concerned mitotic cells (Fig 2B and 2C): in 31 cases, the entire mitotic cell left the sphere (Fig 2B and Expanded Movies 1 and 2), whereas in 4 cases, only one of two daughter cells was budding (Fig 2C). Remarkably, some disseminating cells clearly continued dividing after evading from the sphere (Fig 2C; Expanded Movie 2) and immunofluorescence performed at the end of time lapse microscopy confirmed that budding mitotic cells were not generally engaged in apoptosis, as judged by absence of staining for cleaved caspase 3 (Fig EV3, Expanded Movie 3). When cell cycle progression was arrested at the G1/S transition or in late G2, by addition of thymidine or the CDK1 inhibitor RO3386, respectively, budding of cells from NLP-expressing acini was markedly suppressed (Fig 2D), confirming the dependency of budding on mitosis. We conclude that NLP overexpression specifically triggers the dissemination of mitotic cells from epithelia toward the surrounding matrix.

**Figure 2:**
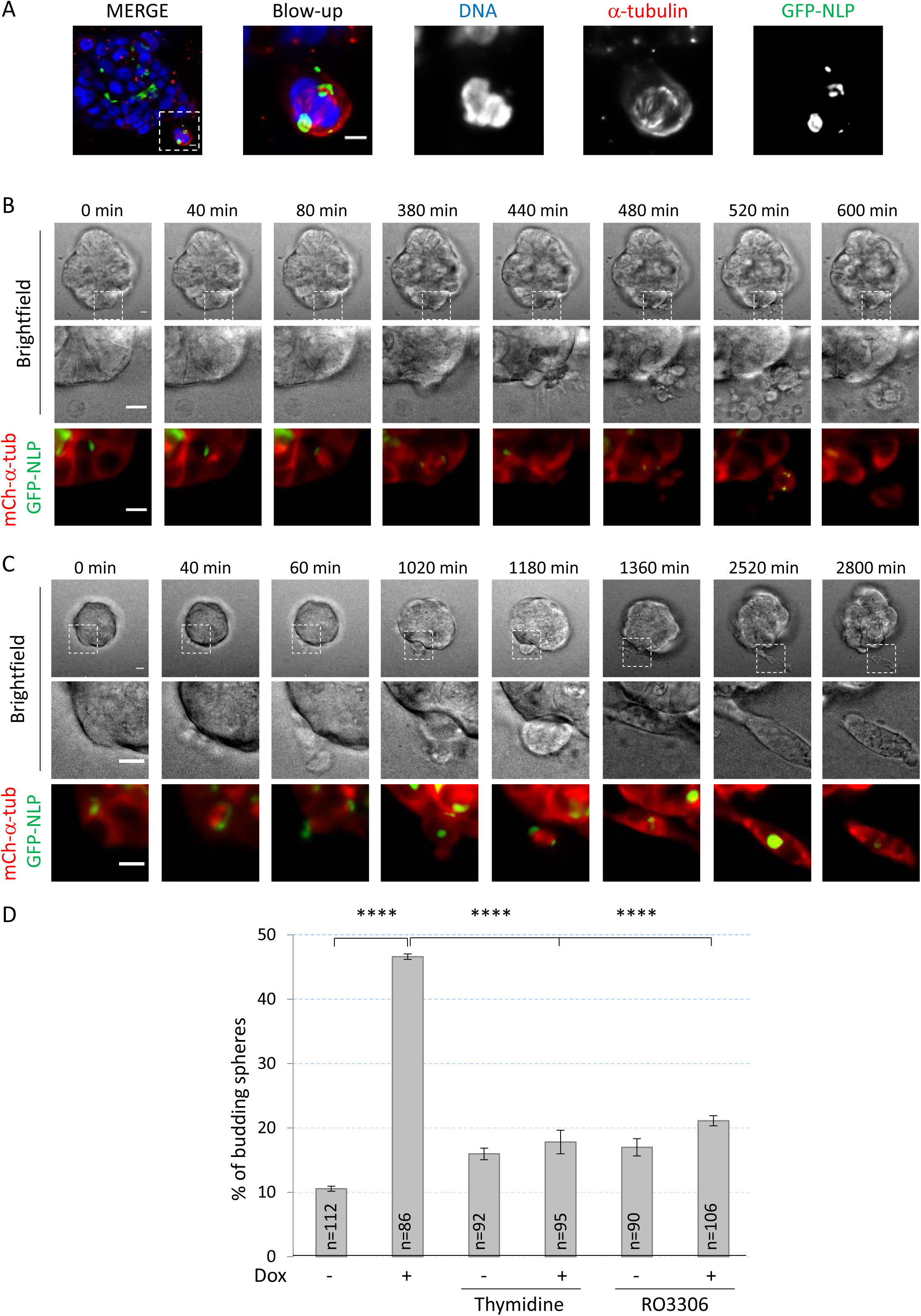
NLP overexpression induces basal budding of living mitotic cells. A. MCF10A acini stained for α-tubulin (red) and DNA (blue) by immunofluorescence. Images show cells that are in mitosis and apparently budding from the parental acini. B-C. Still series from time-lapse experiments showing budding of entire mitotic cell (B) or one of two mitosis-derived daughter cells (C) from MCF10A acini expressing GFP-NLP. Upper panels show brightfield microscopy, middle panels show details of the mitotic budding cells and lower panels the corresponding fluorescence images (GFP-NLP in green and mCherry-α-tubulin in red). Scale bars = 10 µm. Images were acquired every 20 minutes and time stamps are indicated. D. MCF10A acini were treated (+Dox) or not (-Dox) to induce GFP-NLP expression; where indicated, thymidine or RO3306 where concomitantly added to block cells at the G1/S transition or to inhibit CDK1 activity and induce a block in G2 phase, respectively. Fraction of budding acini, with n indicating the number of acini analyzed; error bars indicate ± s.d. of the mean from 3 independent experiments. (****) indicates a P-value < 0.0001, as derived from unpaired, two-tailed Student’s t-test.

### Centrosome aberrations destabilize E-Cadherin junctions

Overexpression of NLP caused disordering of E-Cadherin junctions in both growing MCF10A acini (Schnerch & Nigg, 2016) and MDCK cysts (Fig 3A). As shown by Western blotting, this effect cannot be attributed to altered expression of E-Cadherin (Figure 3B), suggesting that excess NLP interferes with E-Cadherin localization. To quantify the observed mislocalization phenotype, we performed immunofluorescence microscopy on 2D cultures of MDCK cells and calculated an E-Cadherin junction strength index (JSI; see Materials and Methods) (Fig EV4). While already established E-Cadherin junctions within confluent cell layers showed only minor alterations in response to numerical or structural centrosome aberrations (Fig 3C-D, blue bars), E-Cadherin junctions were strongly reduced when transgenes coding for either NLP or PLK4 were induced in exponentially growing cells before these reached confluence (Fig 3C-D, red bars). This suggests that centrosome aberrations interfered with the establishment of E-Cadherin junctions in proliferating cells, consistent with previous data demonstrating that E-Cadherin remodeling occurs during mitosis (Bauer et al., 1998; Ragkousi & Gibson, 2014).

**Figure 3:**
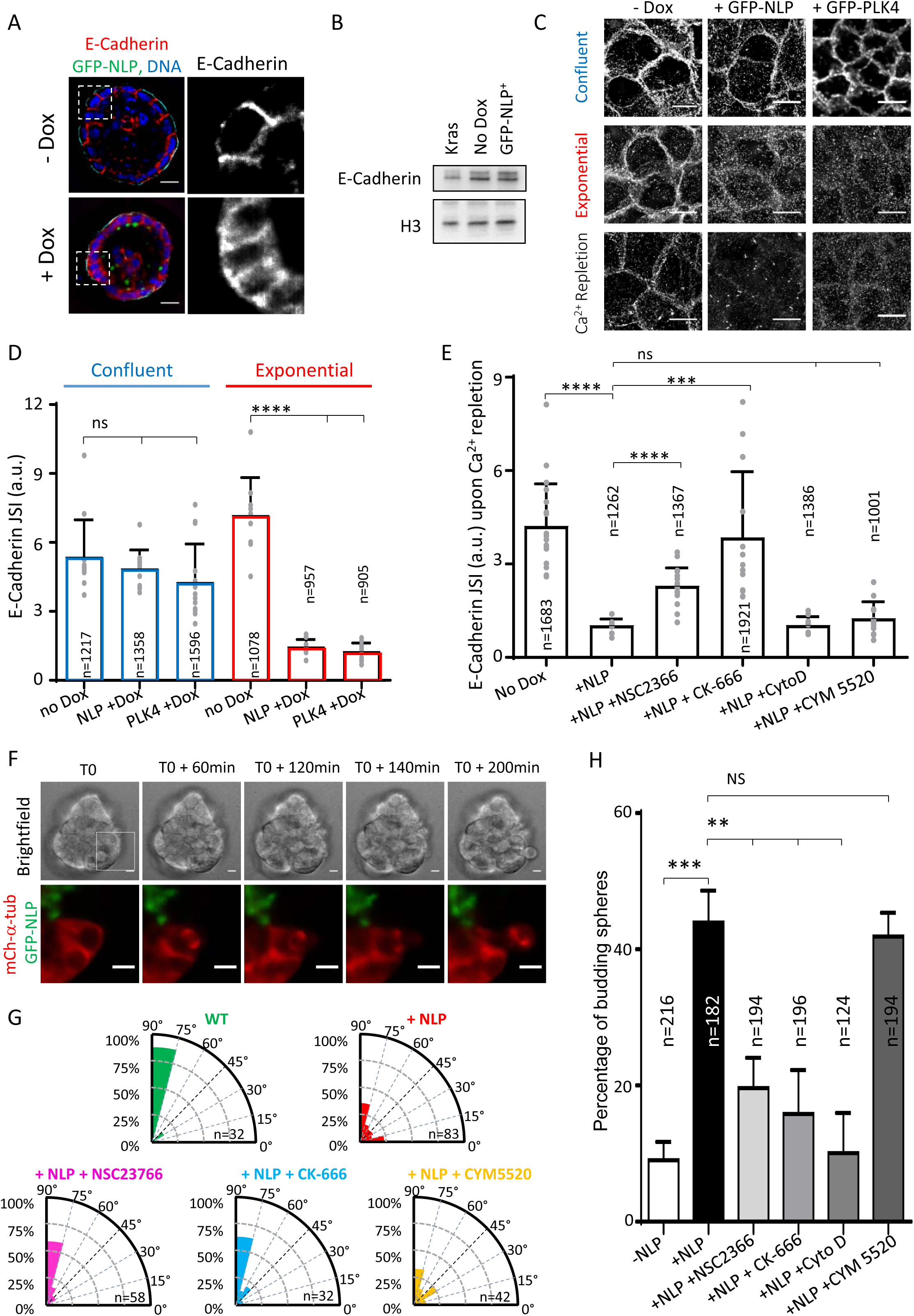
Centrosome aberrations interfere with the establishment of E-Cadherin junctions through Rac1-Arp2/3. A. Representative confocal images showing 8 days old MDCK cysts induced (+Dox) or not (-Dox) to express GFP-NLP during the last 3 days. Cysts were fixed and stained for E-Cadherin (red) and DNA (blue). Note that GFP-NLP overexpression induces a broadening of the E-Cadherin signal, as compared to wild type cysts. Scale bars= 10 µm. B. Total extracts were prepared from MCF10A monolayer cultures overexpressing GFP-NLP (GFP-NLP+), K-Ras (KRas, for control), or no transgene (no DOX) for 3 days, and analyzed by Western blotting using antibodies against E-Cadherin or Histone H3 (loading control). C. Immunofluorescence images show the impact of overexpression of GFP-NLP or GFP-PLK4 on E-Cadherin junctions; transgenes were either induced in confluent MDCK cells (upper panels) or in proliferating cells prior to confluence (middle panels). Also shown are images illustrating the impact of GFP-NLP or GFP-PLK4 on E-Cadherin remodeling induced by calcium (Ca^2+^) repletion (lower panels; for experimental procedure see Fig EV4). All images are z-projected stacks of E-Cadherin staining through the entire height of the cells. Scale bars = 10 µm. D. Histogram represents the E-Cadherin junction strength index (JSI), as defined in Fig EV4, for MDCK cells induced to express GFP-NLP (NLP +Dox), GFP-PLK4 (PLK4 +Dox) or neither transgene (no Dox) during confluence or exponential growth (corresponding to C); Bars represent the means of 3 independent experiments +s.d. and n the numbers of cells analyzed; the values obtained for each field are plotted on the graph. E. Histogram shows the E-Cadherin JSI for calcium repletion experiments performed on confluent cells expressing GFP-NLP (+NLP) or not (-no DOX), and treated with or without the indicated compounds: NSC2366, inhibitor of Rac1; CK-666, inhibitor of Arp2/3; cytochalasin D (CytoD), inhibitor of actin polymerization; CYM5520, an agonist of S1PR2. Bars represent the means of 3 independent experiments +s.d. and n the numbers of cells analyzed; the values obtained for each field are plotted on the graph. F. Still series from time-lapse experiment showing extensive rotation of the mitotic spindle during budding from an MCF10A acinus stably expressing mCherry α-tubulin (red) and induced to express GFP-NLP (green). Upper panels show brightfield images and lower panels the corresponding fluorescence images (see Expanded Movie 4). Scale bars = 10 εm G. Radial histograms illustrate the distributions of the acute angle between the mitotic spindle axis and the radius of the cyst for the different conditions on fixed samples. n represent the number of mitoses analyzed. Note that data for NLP^+^ cysts include both GFP-NLP^−^ and GFP-NLP^+^ mitotic cells, as results for these two subclasses were virtually indistinguishable. This confirms that spindles rotate in both GFP-NLP^+^ and GFP-NLP^−^ mitotic cells budding from NLP^+^ cysts (as illustrated in panel F). H. Fraction of budding acini in response to the indicated treatments. Bars represent means +s.d. and n the number of acini from 3 independent experiments. P-values were derived from unpaired, two-tailed Student’s t-test. ns indicates not significant; (*), (**), (***) and (****) indicate P-values of <0.05, <0.005, <0.0005 and <0.0001, respectively.

To obtain information about the E-Cadherin regulatory pathway perturbed by NLP overexpression, we next applied an established calcium repletion procedure (Le et al., 1999) to confluent MDCK cells harboring normal or aberrant centrosomes (Fig EV4). While control cells readily re-established E-Cadherin junctions upon calcium repletion, cells harboring NLP-induced structural centrosome aberrations were unable to do so (Fig 3C, bottom row). Numerical centrosome aberrations triggered by PLK4 overexpression also interfered with junction formation (Fig 3C, bottom row), confirming previous observations (Godinho et al., 2014). An excess of growing microtubules was previously shown to destabilize E-Cadherin junctions by altering the cortical actin network *via* over-activation of the Rac1-Arp2/3 effectors (Akhtar & Hotchin, 2001; Chu et al., 2004; Godinho et al., 2014; Waterman-Storer et al., 1999; Xue et al., 2013). Thus, we asked whether NLP overexpression interfered with this pathway. Indeed, partial reduction of Rac1 or Arp2/3 activity, using the respective inhibitors NSC-23766 and CK-666, substantially prevented the NLP-induced inhibition of E-Cadherin junction formation (Fig 3E). In contrast, no rescue was seen with cytochalasin D, an inhibitor of actin polymerization, or with CYM5520, an agonist of the sphingosine 1 receptor 2 (S1PR2) pathway implicated in basal extrusion (Slattum et al., 2014, Hendley et al. 2016) (Fig 3E) (see Discussion). This argues that NLP overexpression impairs E-Cadherin junctions via increased stabilization of microtubules, which then results in excessive activation of the Rac1-Arp2/3 pathway. In support of this conclusion, it was previously shown that NLP overexpression results in the accumulation of microtubules at structurally enlarged centrosomes (Casenghi et al., 2003; Schnerch & Nigg, 2016). Moreover, these microtubules are trapped *via* their minus-ends, as shown by staining for CAMSAP-2 (Akhmanova & Hoogenraad, 2015; Jiang et al., 2014) (Fig EV5), and they display enhanced stability, as revealed by staining with antibodies against de-tyrosinated α-tubulin (Kerr et al., 2015; Schulze et al., 1987; Webster et al., 1987) (Fig EV5).

To confirm that mitotic cell-cell contacts are impaired in response to NLP overexpression, we used time lapse microscopy to monitor junction strength in 2D MCF10A cultures. As summarized in Fig EV6, we found that NLP overexpression reduced the time between the metaphase onset and the loss of contact between a dividing cell and adjacent cells. This contributes to explain the selective budding of mitotic cells from 3D acini.

E-Cadherin junctions are crucial for the preservation of the orientation of cell divisions within epithelial tissues (Gloerich et al., 2017; Ragkousi & Gibson, 2014). In line with this notion, we observed marked spindle rotation in budding mitotic cells (Fig 3F, Expanded Movie 4). To quantify this rotation phenotype, we also measured spindle orientation on fixed samples. In wild-type MDCK cysts, the mitotic spindles were preferentially oriented orthogonally to the apico-basal cell axis, as expected (Fig 3G). In contrast, spindle orientation was randomized in cysts harboring NLP-induced centrosome aberrations (Fig 3G), corroborating the weakening of interactions between these latter mitotic cells and their neighbors. Partial inhibition of Rac1 (NSC23766) or Arp2/3 (CK-666) in these cysts restored wild-type spindle orientation (Fig 3G), confirming that the Rac1-Arp2/3 pathway regulates E-Cadherin junctions. In contrast, CYM5520, the compound inhibiting basal extrusion (Hendley et al., 2016; Slattum & Rosenblatt, 2014), did not rescue the phenotype (Fig 3E-G) (see Discussion). We emphasize that the ability of these three inhibitors to restore E-Cadherin junctions and spindle orientation was correlated with their ability to prevent NLP-induced budding (Fig 3H).

Taken together, the above results show that centrosome aberrations interfere with E-Cadherin junctions. Importantly, however, this E-Cadherin phenotype was elicited by both structural and numerical centrosome aberrations (Fig 3A; see also (Godinho et al., 2014)), yet only NLP overexpression induced budding, while PLK4 overexpression did not (Figure 1B). Conversely, even though Cytochalasin D prevented NLP-induced budding, it did not restore the establishment of E-Cadherin junctions (Fig 3E-H). Taken together, these data demonstrate that weakening of mitotic E-Cadherin junctions is necessary but not sufficient to explain the budding of mitotic cells in response to NLP-induced centrosome aberrations, implying the involvement of at least one additional mechanism.

### Budding represents a non-cell autonomous process

Because not all cells express the GFP-NLP transgene product upon induction (Fig EV 7 and 8), we asked whether the extent of budding could be correlated to the percentage of GFP-NLP^+^ cells within mosaic acini. After estimating the proportion of GFP-NLP^+^ cells per acinus (for details see Materials and Methods), these were classified into 4 classes differing in the proportion of GFPNLP^+^ cells, and the frequency of budding was determined for each class (Fig 4A). This analysis revealed that the frequency of budding is not proportional to the percentage of GFP-NLP^+^ cells. Instead, extensive budding is triggered once the proportion of GFP-NLP^+^ cells exceeds a threshold (> 50%), strongly suggesting that budding requires multicellular cooperation within the epithelium. Considering that the parental mosaic acini displayed 50 to 100% of GFP-NLP^+^ cells, it is remarkable that less than 50 % of all budding cells expressed detectable levels of GFP-NLP (illustrated in Fig 4A-B and Fig EV3). This demonstrates that budding reflects a non-cell autonomous mechanism and indicates that GFP-NLP^−^ cells are preferentially squeezed out over GFP-NLP^+^ cells. This may establish a positive feed-back loop that progressively enriches the epithelium with GFP-NLP^+^ cells and favors the dissemination of mitotic cells that do not themselves harbor centrosome aberrations.

**Figure 4:**
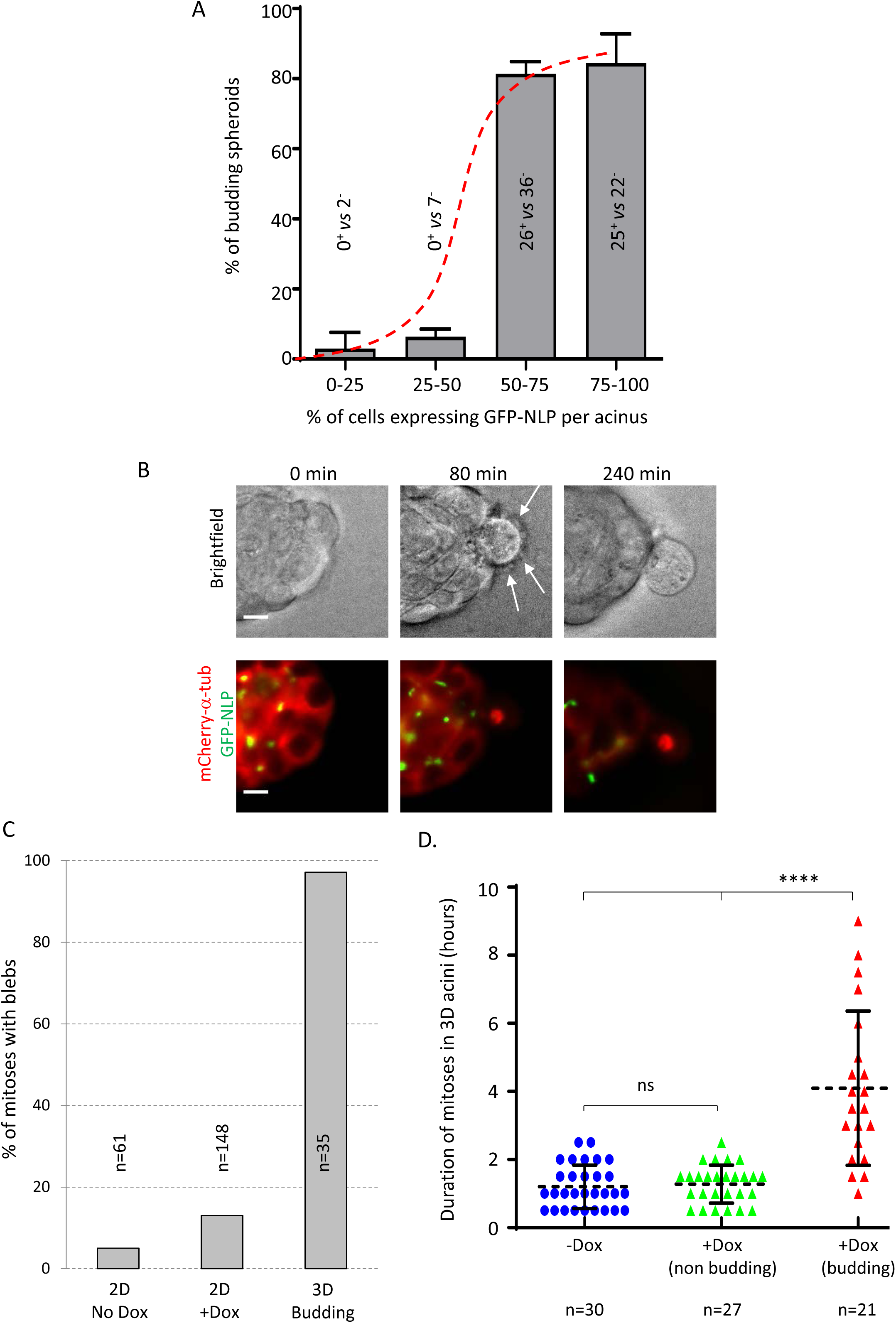
Budding of mitotic cells requires multicellular cooperation and reflects a non-cellautonomous process. A. Fraction of budding acini as a function of the estimated percentage of GFP-NLP^+^ cells. Acini were grouped into 4 classes, depending on the percentage of GFP-NLP^+^ cells within each acinus. As illustrated by the sigmoidal curve (red dashed curve), the frequency of budding is not proportional to the percentage of GFP-NLP^+^ cells within each group, but instead requires a threshold (> 50%) above which budding occurs with high frequency. All budding cells were also analyzed for expression of GFP-NLP and the numbers of GFP-NLP^+^ versus GFP-NLP^−^ budding cells are indicated for each class. This revealed 0% of GFP-NLP^+^ budding cells in the 0-25% (n=2) and 25-50% classes (n=7), 46% in the 50-75% class (n=62) and 53% in the 75-100% class (n=47). Bars represent the means of 4 independent experiments +s.d. of 293 spheres analyzed. B. Representative stills from time lapse recordings show a mitotic cell budding from an MCF10A acinus stably expressing mCherry α-tubulin (red) and induced to express GFP-NLP (green). Upper panels show brightfield images and lower panels the corresponding fluorescence images; arrows point to membrane blebs. Images were acquired every 20 minutes and time stamps are indicated; scale bars = 10 µm. C. Fraction of mitotic cells associated with extensive blebbing in different MCF10A populations observed by time lapse for 2 days. The graph compares MCF10A cells cultured in a 2D monolayers without (2D No Dox) or after induction of GFP-NLP expression (2D +Dox) and budding cells from 3D acini (3D Budding). Bars are means of 2 independent experiments for the 2D cultures and of 16 independent experiments for the 3D acini; n represents the number of mitoses analyzed. D. Scatter plot shows the mitotic duration, determined from time lapse experiments, of cells dividing within MCF10A acini. The graph compares mitoses in acini without GFP-NLP induction (-Dox, blue circles) and mitoses within acini expressing GFP-NLP (+Dox) that either undergo budding (budding, red triangles) or not (non-budding, green triangles). Error bars represent ±s.d. of the means and n shows the numbers of mitoses analyzed. P-values were derived from unpaired, two-tailed Student’s t-test with Welch’s correction; (****) indicates a P-value < 0.0001 and ns means not significant.

Budding mitotic cells displayed two striking characteristics. First, almost all budding cells monitored by time-lapse microscopy displayed extensive membrane blebbing (Fig 4C). This blebbing was also seen in budding cells that lacked detectable GFP-NLP (Fig 4B, Expanded Movies 1-3), indicating that it is not a cell autonomous consequence of NLP over-expression. Corroborating this conclusion, NLP overexpression induced only a minor increase in mitotic blebbing in 2D cultures (13% *vs* 5% for control cells), in stark contrast to the blebbing cells seen in 97% of cells budding from acini (see Fig 4C). Second, budding cells showed markedly extended M phase durations and typically remained in metaphase for several hours (Fig 2B and 4D). In contrast, non-budding mitoses within acini overexpressing NLP lasted approximately 1 hour, similar to the duration of mitoses in non-induced controls or 2D cultures (Fig 4D; Fig EV8). This demonstrates that both blebbing and prolonged mitoses are related to budding rather than NLP overexpression *per se*.

### Budding is governed by biomechanical properties of epithelia

Both blebbing and delayed mitotic progression have recently been observed in confined mitotic cells exposed to increasing pressure (Cattin et al., 2015). This led us to postulate that cells harboring structural centrosome aberrations might display altered biomechanical properties. If such cells were to display increased stiffness, epithelia containing a sufficient proportion of GFP-NLP^+^ cells might confine mitotic cells and eventually squeeze them out. To explore this hypothesis, we used atomic force microscopy (AFM) to map the mechanical stiffness of various cell populations. Indeed, when compared to cells overexpressing either PLK4 or CEP68 within a confluent MDCK monolayer, interphase cells overexpressing NLP showed a significant increase in cellular stiffness (Fig 5A, Fig EV9a). Furthermore, probing the basal surface of MDCK 3D cysts revealed a similar increase in cellular stiffness in response to NLP overexpression (Fig 5B). For comparison, we also measured cellular stiffness within confluent MDCK monolayers after application of different drugs acting on the cytoskeleton (Fig 5C). Treatment with taxol, a stabilizer of microtubules, also increased stiffness, in line with previous results (Kerr et al., 2015). In wild-type cells, the taxol-induced stiffness was comparable to that seen upon overexpression of NLP, but in GFP-NLP^+^ cells, which already contain stabilized microtubules (Fig EV5), taxol produced only a minor additional effect. Conversely, inhibition of microtubule polymerization by nocodazol did not affect wild-type cells and slightly reduced cellular stiffness in GFP-NLP^+^ cells (Fig 5C). The relatively minor but significant effect falls in line with immunofluorescence experiments showing that nocodazol treatment caused only partial depolymerization of microtubules in confluent interphase cells (Fig EV9B; see also (Pepperkok et al., 1990)). Finally, inhibition of actin polymerization by cytochalasin D reduced the stiffness of both control cells and GFP-NLP^+^ cells (Fig 5C), confirming that actin exerts a major contribution to cellular stiffness (Bruckner & Janshoff, 2015; Fletcher & Mullins, 2010). Taken together, these results support the notion that NLP-induced centrosomal aberrations increase cellular stiffness by stabilizing microtubules, which in turn influences the actin cytoskeleton.

**Figure 5:**
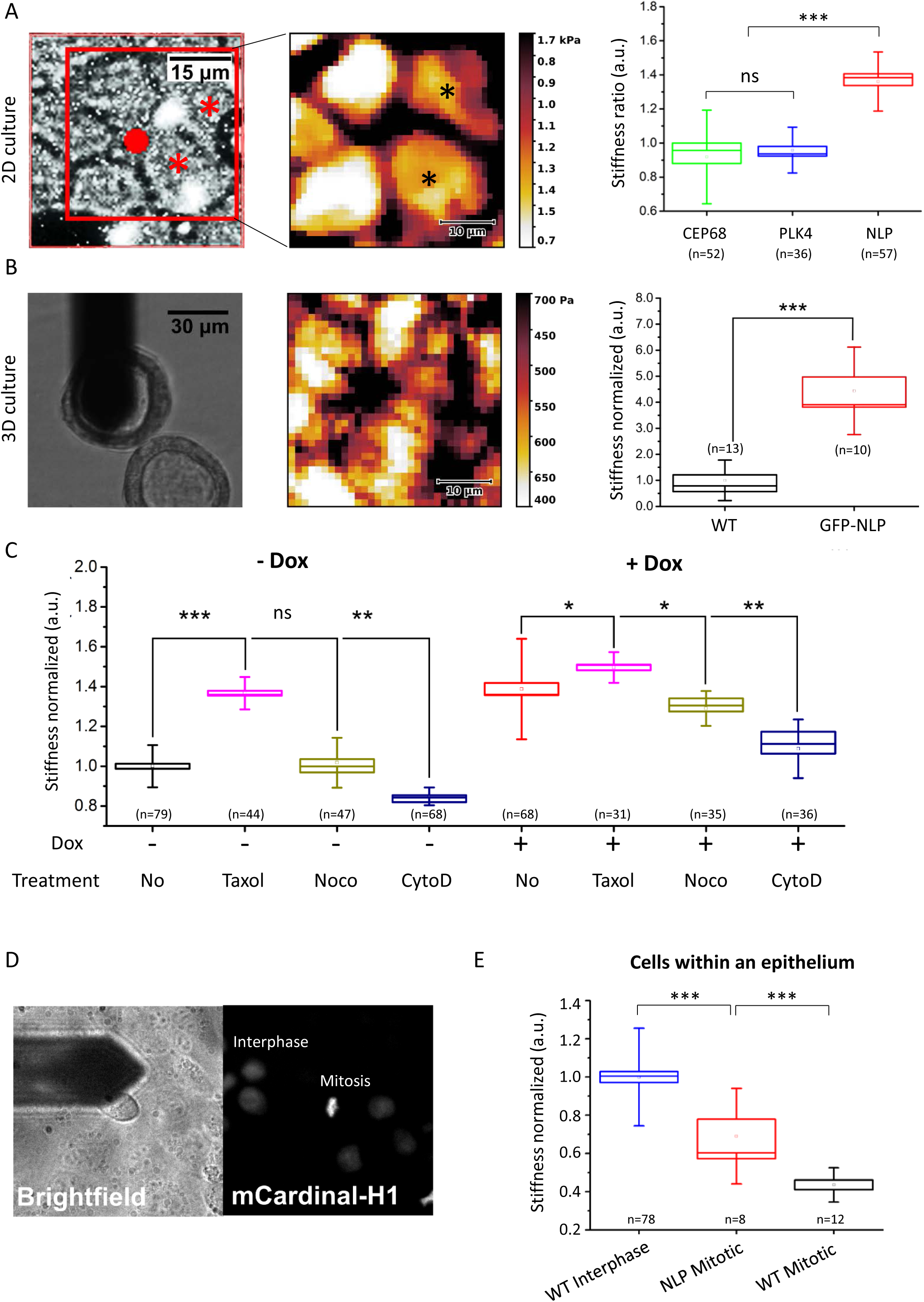
Structural but not numerical centrosome aberrations increase stiffness of epithelial cells. A. AFM performed on MDCK cells cultured in 2D. (Left) Example of epifluorescence microscopy image; cells harboring structural centrosome aberrations (GFP-NLP) are marked (asterisks). The scan area (red square) and position of the AFM probe (red dot) are monitored in the brightfield signal for each experiment. (Middle) Corresponding stiffness map visualizes local nanomechanical heterogeneities; cells overexpressing GFP-NLP, as visualized by epifluorescence microscopy, are marked by asterisks. (Right) Stiffness ratios compare cells overexpressing NLP, PLK4 or CEP68 with neighboring non-expressing cells (WT); NLP-over-expressing cells are stiffer than WT cells while cells over-expressing either PLK4 or CEP68 exhibit mechanical phenotypes similar to WT cells. B. AFM performed on 3D MDCK cysts. (Left) brightfield image showing the AFM probe positioned above the isolated cyst. (Middle) stiffness map visualizes basal surface of individual cells within top region of 3D cysts. (Right) quantitative analysis confirms that NLP over-expression induces cell stiffening in 3D. C. Drug-induced changes of cytoskeletal structures alter cellular stiffness of WT (-Dox) or GFPNLP^+^ (+Dox) MDCK cells in 2D. D. Cell cycle dependence of cellular stiffness. Position of the AFM probe is monitored in the brightfield signal while mCardinal-H1 signal detection by epifluorescence allows identification of mitoses and interphase cells. Images are 90 × 90 µm. E. Quantitative AFM analysis of confluent MDCK cells in interphase or in mitosis, in comparison to surrounding cells in interphase that do not express GFP-NLP (WT). n indicates the number of analyzed cells. Box plots show the mean (square) and median (line); whiskers are s.d. and the box is s.e.m. Statistical significance was tested using a Mann-Whitney test. (*), (**) and (***) indicate P-value <0.05, <0.01 and <0.005 respectively.

As budding selectively involves mitotic cells, we next compared the stiffness of interphase and mitotic cells. Specifically, we analyzed stiffness in near-confluent monolayers of MDCK cells expressing mCardinal-H1 to allow monitoring of cell cycle phases (Fig 5D). Interphase cells were generally stiffer than mitotic cells and overexpression of NLP increased the stiffness of the latter (Fig 5E). These results fall in line with previous *in situ* observations, suggesting that the presence of soft cells in tumor biopsies correlates with metastatic spreading (Lekka et al., 2012; Plodinec et al., 2012; Swaminathan et al., 2011), and they suggest that the higher stiffness of GFP-NLP^+^ mitotic cells contributes to explain the preferential budding of wild-type mitotic cells (Fig 4B, (Cadart et al., 2014)). However, these results seemed to conflict with earlier data showing that cells become stiffer when entering mitosis (Chugh et al., 2017; Kunda et al., 2008; Stewart et al., 2011). We suspected that this difference might relate to the fact that previous measurements were performed on isolated single cells lacking cell-cell contact. Indeed, we could readily confirm that isolated MDCK cells going through mitosis display a higher stiffness than interphase cells (Fig EV 9C). These data demonstrate that cellular stiffness is drastically influenced by physical constraints, notably the confinement of cells within an epithelium (Saw et al., 2017). Collectively, our data lead us to propose that centrosome aberrations trigger not only cytoskeletal remodeling but also heterogeneity in the biomechanical properties of epithelia, which then results in the selective budding of mitotic cells through a non-cell autonomous process (see schematic model in Fig EV10).

## DISCUSSION

Centrosome aberrations are common in human tumors (Chan, 2011; Guo et al., 2007; Lingle et al., 1998), but their role in carcinogenesis remains subject to intense debate (Godinho & Pellman, 2014; Gonczy, 2015; Nigg, 2002; Raff & Basto, 2017). Studies in cells and animals support the view that centrosome amplification induces aneuploidy through chromosome mis-segregation (Ganem et al., 2009; Levine et al., 2017), but other possible contributions to cancer development have also been proposed (Basto et al., 2008; Coelho et al., 2015; Godinho et al., 2014; Kazazian et al., 2017; Sercin et al., 2016). Yet, a conundrum persists in that centrosome aberrations are *a priori* expected to impair the viability of those tumor cell subpopulations that harbor these aberrations. Thus, the functional significance of centrosome aberrations in human tumors remains difficult to ascertain. Our study identifies a novel mechanism through which cells harboring centrosome aberrations may contribute to promote an invasive phenotype through a non-cell autonomous process, thereby offering a solution to the above conundrum. Specifically, we propose a model with the potential to explain how centrosome aberrations could contribute to metastasis, without the disseminating cells carrying these deleterious alterations (EV10).

We show that structural centrosome aberrations, induced by overexpression of NLP (Casenghi et al., 2003; Schnerch & Nigg, 2016), trigger the selective budding of mitotic cells from 3D epithelial acini. Furthermore, we identify two complementary mechanisms supporting this cell dissemination: first, increased microtubule stability in GFP-NLP^+^ cells causes a weakening of E-Cadherin junctions between mitotic cells and their neighbors. Second, cytoskeletal aberrations induce heterogeneity in the biomechanical properties (stiffness) of cells, causing extensive blebbing and marked delays in mitotic progression, indicative of confinement (Cattin et al., 2015; Sorce et al., 2015). Consequently, when epithelia contain cells whose stiffness is increased by centrosome aberrations, soft mitotic cells are selectively squeezed out. Considering that mitotic cells devoid of centrosome aberrations are softer than GFP-NLP^+^ mitotic cells, the former are expected to offer reduced resistance to extrusion forces (Cadart et al., 2014; Sorce et al., 2015) and hence bud preferentially, as observed.

In recent years, a delamination process known as basal epithelial cell extrusion has attracted increasing attention (Marshall et al., 2011; Slattum et al., 2009; Slattum & Rosenblatt, 2014). This process contributes to remove dying or unwanted cells from epithelia (Eisenhoffer et al., 2012; Slattum et al., 2014) and has also been proposed to play a role in tumor cell invasion (Slattum & Rosenblatt, 2014). However, although the budding mechanism described here can formally be considered as a form of basal extrusion, we emphasize that the underlying mechanism differs in several fundamental aspects from the basal extrusion phenomenon pioneered by Rosenblatt and coworkers. First, basal extrusion of dying cells is based on constriction of an actomyosin ring (Marshall et al., 2011; Slattum et al., 2014; Slattum et al., 2009) and considering that apical activation of actomyosin contractility is incompatible with mitosis (Grosshans & Wieschaus, 2000; Mata et al., 2000; Seher & Leptin, 2000), such a mechanism is not consistent with the budding of mitotic cells. Second, basal extrusion is inhibited by CYM5520, an agonist of S1PR2 (Hendley et al., 2016), while we show here that this compound does not prevent budding (Fig 3). Finally, while basal extrusion is a cell-autonomous phenomenon (Slattum et al., 2014), budding represents a non-cell-autonomous phenomenon based on multicellular cooperation. Hence, cell dissemination by budding is mechanistically distinct from basal extrusion.

NLP is frequently overexpressed in different human cancers (Qu et al., 2008; Shao et al., 2010; Yu et al., 2009) and reported to confer resistance to paclitaxel in breast cancer (Zhao et al., 2012). Moreover, NLP-induced centrosome aberrations in 2D and 3D culture models recruit γtubulin and centrin (Casenghi et al., 2003; Schnerch & Nigg, 2016), highly reminiscent of the aberrations seen in human tumors (Guo et al., 2007; Lingle et al., 1998; Salisbury et al., 2004). Accordingly, we used NLP to study the impact of structural centrosome aberrations on epithelial architecture and invasiveness. In future, it will be interesting to determine whether other centrosomal proteins affecting the dynamics of the microtubule network (Delaval & Doxsey, 2010; Fogeron et al., 2013) trigger a similar invasive phenotype. It is tempting to speculate that any cellular alteration able to exert comparable effects on both E-Cadherin junctions and the biomechanical properties of epithelia may trigger the dissemination of mitotic cells through a similar non-cell-autonomous mechanism.

The dissemination of individual cells or cell clusters, accompanied by complete or partial epithelial-mesenchymal transition (Nieto et al., 2016), is considered a first key step in metastasis (Aceto et al., 2014; Lambert et al., 2017). Moreover, the importance of short-range dispersal of individual tumor cells is well recognized (Waclaw et al., 2015). A complete understanding of cell dissemination will ultimately require a combination of intra-vital imaging (Alexander et al., 2013; Paul et al., 2017) and simulation of invasive processes using defined *in vitro* systems (Discher et al., 2017; Shamir & Ewald, 2014). Results obtained with established 2D and 3D culture models lead us to propose a novel mechanism through which centrosome aberrations could trigger the dissemination of incipient tumor cells. Extrapolating to an *in vivo* situation, our findings have several implications. First, they bear on the question of when disseminating cancer cells first arise (Ghajar & Bissell, 2016). Considering that centrosome aberrations can be observed already in premalignant lesions, the mechanism proposed here would allow dissemination of cells with metastatic potential from very early tumors, in line with recent proposals (Harper et al., 2016; Hosseini et al., 2016). Second, we conclude that cytoskeletal aberrations affecting tissue architecture need not necessarily occur within the same cells that harbor oncogenic mutations, offering a new explanation for how centrosome aberrations could contribute to aggressive cancer development in spite of being *a priori* deleterious. Third, the non-cell-autonomous nature of the observed process implies that aberrations conferring metastatic properties may not necessarily be detectable within the disseminating cancer cells themselves, implying that drivers of metastasis may escape detection by genetic methods comparing metastatic cells with primary tumor cells. Collectively, our data contribute to focus attention on the microenvironment surrounding tumors cells (Bissell & Hines, 2011; Tabassum & Polyak, 2015) and on the biomechanical properties of tumor tissues (Lee et al., 2012; Plodinec et al., 2012; Swaminathan et al., 2011). In particular, our data support previous observations suggesting that metastatic spreading correlates with the presence of low stiffness cells within tumor biopsies (Plodinec et al., 2012; Swaminathan et al., 2011). Finally, our findings may also have implications for normal development. In particular, it will be interesting to explore whether differences in stiffness could contribute to trigger developmentally controlled epithelial invaginations for which mitotic progression appears to be a pre-requisite (Kondo & Hayashi, 2013).

## MATERIALS AND METHODS

### Generation of expression constructs and cell lines

cDNA encoding mCherry-α-tubulin, mCardinal-H1 and dTomato-VE-Cadherin were PCR-amplified from pmCherry_α_tubulin_IRES_puro2 (kindly provided by Daniel Gerlich (Steigemann et al., 2009)), mCardinal-H1-10 (a gift from Michael Davidson (Chu et al., 2014) (Addgene plasmid # 56161)), and tdTomato-VE-Cadherin-N-10 (a gift from Michael Davidson (Addgene plasmid # 58142) respectively and subsequently ligated into pMXs-IRES-Blasticidin (Cell Biolabs Inc.).

MCF10A ecoR cell lines allowing doxycycline-inducible expression of EGFP-NLP, EGFP-CEP68 and EGFP-PLK4 were described previously (Schnerch & Nigg, 2016). Doxycycline-inducible MDCK II cells were generated using the same two-step transduction strategy and enriched by antibiotic selection using hygromycin at 400 μg.mL^−1^ and puromycin at 2 μg.mL^−1^. Inducible MDCK and MCF10A cells stably expressing mCherry-α-tubulin, dTomato-E-Cadherin and m-Cardinal H1 were generated by retroviral transduction and sorted by flow cytometry using BD FACSAria IIIu cell sorter (FACS Core Facility of Biozentrum). MCF10A ecoR cells stably expressing K-Ras were obtained by ecotropic retroviral transduction with pBabe K-Ras 12V (Addgene plasmid #12544, gift from Channing Der (Khosravi-Far et al., 1996)).

### Cell Culture

MCF10A ecoR cells (a kind gift from Tilman Brummer; University of Freiburg) were grown as described previously (Debnath et al., 2003). Briefly, MCF10A cells were grown in DMEM:F12 (Sigma-Aldrich, St. Louis, MO, USA) supplemented with 10 μg.ml^−1^ insulin (Sigma-Aldrich, St. Louis, MO, USA), 0.5μg.ml^−1^ hydrocortisone (Sigma-Aldrich, St. Louis, MO, USA), 100 ng.ml^−1^ cholera toxin (Sigma-Aldrich, St. Louis, MO, USA), and penicillin/streptomycin (Life technologies, Carlsbad, CA, USA), 2% or 5% horse serum (Life technologies, Carlsbad, CA, USA) and 5 ng.ml^−1^ or 20 ng.ml^−1^ epidermal growth factor (Peprotech, London, UK) for 3D and 2D culture respectively. MDCK II cells (a kind gift from Inke Naethke, University of Dundee, UK) were grown in Minimum Essential Medium Eagle (Sigma-Aldrich, St. Louis, MO, USA) supplemented with 10% fetal calf serum (GE Healthcare, Chicago, Illinois, USA) and Penicillin Streptomycin (Life technologies, Carlsbad, CA, USA). Phoenix cells and HEK293T cells (provided by Stefan Zimmermann and Ralph Wäsch; University Medical Center Freiburg) were grown in DMEM medium (Life technologies, Carlsbad, CA, USA) supplemented with 10% fetal calf serum (GE Healthcare, Chicago, Illinois, USA), sodium pyruvate (Life technologies, Carlsbad, CA, USA) and Penicillin Streptomycin (Life technologies, Carlsbad, CA, USA). Tissue cultures were routinely tested for mycoplasma contamination by PCR using growth medium from high density cultures as a template. MCF10A acini and MDCK II cysts were generated by plating single cell solutions onto beds of pure Matrigel (356231, Corning) or Matrigel enriched by collagen I (1.6 mg.mL^−1^, Life technologies, Carlsbad, CA, USA), as described previously (Debnath et al., 2003; Godinho et al., 2014). To monitor invadopodia formation, MCF10A cells were seeded in Collagen I-enriched Matrigel and transgene expression was induced by doxycycline the day following seeding.

### Compounds

Expression of transgenes coding for EGFP-tagged centrosomal proteins was initiated by addition of 2.5 μg.ml^−1^ of doxycycline. Inhibitors used in 3D acini experiments and for determination of ECadherin JSIs were used at the following concentrations: 25 μM NSC23766 (Sigma Aldrich MO, USA), 50 μM CK-666 (Sigma Aldrich MO, USA), 2.5 μM cytochalasin D (Sigma Aldrich MO, USA), 5μM RO-3306 (Merck Millipore Darmstadt Germany), 2 mM Thymidine (Sigma Aldrich MO, USA). For rescue experiments, doxycycline and drugs were added simultaneously to medium.

### Fluorescence microscopy

2D monolayers of MDCK II or MCF10A cells were grown and processed for immunolabeling in Ibidi 8 well microscopy slides, and 3D acini derived from MCF10A or MDCK cells were grown in 8 chambers slides (354108, Falcon Corning) and processed for immunostaining essentially as described previously (Debnath et al., 2003; Schnerch & Nigg, 2016). Briefly, cells and cysts were fixed using 2% formalin or 4% paraformaldehyde in PBS for 15 minutes at room temperature. Cells were permeabilized using PBS 0.5% Triton X-100 for 5 minutes and blocked in 2% BSA in PBS for 30 minutes and antibodies diluted in PBS-0.5%Tween-2%BSA. Primary antibodies used were anti-α-tubulin (T9026, Sigma, St. Louis, Mo, USA), anti-cleaved-caspase 3 (D175; 9661, Cell signalling Technology) and anti-E-Cadherin (610182, BD, Franklin Lakes, NJ, USA). F-actin fibres were stained using AlexaFluor 647-linked phalloidin at 33 nM (A22287, Life Technologies, Carlsbad CA, USA) and DNA stained with 1 μg.mL^−1^ DAPI. Secondary antibodies were AlexaFluor647 goat anti-mouse (A21236), AlexaFluor568 goat anti-mouse (A11004) and AlexaFluor568 donkey anti-rabbit (A10042) (all from Life Technologies, Carlsbad CA, USA). 3D acini were finally mounted in ProLong Antifade mounting medium (Molecular Probes) and analyzed rapidly.

Confocal images were acquired using a Leica SP5-II-MATRIX point scanning confocal microscope equipped with a 20x/0.70 HCX Plan Apo CS air objective and a 63x/1.40-1.60 HCX Plan Apo lambda blue oil immersion objective. 405nm diode laser light was applied for DAPI staining, 488nm Argon laser light for visualization of GFP, 561nm DPSS laser light for visualization of AlexaFluor568 stainings and 633nm HeNe laser light for visualization of AlexaFluor647 stainings. Image analyses and final adjustments of confocal images taken in the sphere equator planes were carried out in Omero 5.1.2. Image analyses, 3D reconstructions and final adjustments of confocal z-stacks (spacing 0.3 μm between confocal planes) were carried out in Imaris 8.1.2.

For live cell imaging of 3D cultured acini, single cells were seeded in Matrigel in 8 wells slides and processed as described (Debnath et al., 2003) for 5 days until the beginning of the time lapse. The entire acini were filmed by taking stacks of pictured spaced of 0.37 μm. Gridded bottom slides (80827 or 80826-G500, Ibidi Germany) were used when previously recorded acini had to be identified for subsequent immunofluorescence analyses. For time lapse microscopy on 2D monolayers, cells were grown on collagen IV coated ibidi 8 wells slides (80822, Ibidi, Germany) and GFP-NLP expression was induced 48 hours prior to the onset of recording.

All live-cell imaging experiments were carried out using a FEI MORE wide-field system (FEI Munich, Graefelfing, Germany) equipped with a 40x/0.95 U Plan S Apo air objective. For visualization of EGFP- and mCherry-signals, LEDs, combined with a quad bandpass filter, were used as a light source for trans-illumination at 515/18 and 595/19 nm, whereas dTomato- and mCardinal signals were distinguished using single band pass filters at 590 nm and 685 nm, respectively. Pictures acquisition was performed at 37°C, 5% CO_2_ and >70% air humidity. Image analyses were carried out using Image J or Imaris 8.1.2. following deconvolution using Huygens Remote manager 3.3.0-rc9.

### Determination of E-Cadherin junction strength index (JSI)

MDCK cells were stained for DNA (DAPI) and E-Cadherin, and z-stacks (spacing 0.2 μm) of the whole epithelium within the imaged field were imaged (around 70 planes). Images were then processed for deconvolution using Huygens Remote manager 3.3.0-rc9 and z-projections of the maximum intensities of the whole stacks were performed using Fiji software. E-Cadherin signals were processed to determine an E-Cadherin JSI using Icy software (http://icy.bioimageanalysis.org/). Briefly, E-Cadherin signals were segmented into 3 classes, termed background, junctions and speckles/aggregates, by using k-means to define two thresholds. A mask was then created to identify the junction and a median filter with half size 2 used to remove noise. This mask was used to determine the entropy and the area of the signal using Icy’s ROI statistics. E-Cadherin JSI was finally defined as the ratio of area to entropy of signal per nucleus.

### Mitotic spindle angle measurements

3D reconstructions from z-stack images (spacing 0.2 μm) of MDCK cysts stained for α-tubulin and DNA and were performed using Imaris 8.1.2. and the acute angles between the spindle axes and the radius of the cysts (hitting the center of the mitotic spindle); in all cases, spheres were rotated to ensure positioning the mitotic spindle within the equatorial plane of the cyst.

### Flow cytometry

Cells were trypsinized, washed twice with PBS, fixed with PFA 4% for 10 minutes at room temperature (RT), permeabilized for 2 minutes with 0.2% Triton X-100 and then treated with 50µg.ml^−1^ RNAse A (Sigma, R6513) and stained with 25µg.ml^−1^ propidium iodide (PI) for 30 minutes at 37°C. Cells were analyzed with a BD FACS Canto II cytometer and an approximate evaluation of the different cell cycle populations were obtained using the software FlowJo (Tree Star Inc, Ashland, Oregon, U.S.A.).

### Estimation of the percentage of cells GFP-NLP ^+^ within a sphere

The numbers of GFP-NLP^+^ CRBs (centrosome related bodies; Schnerch and Nigg 2016) and nuclei (stained with DAPI) within a given acinus were determined using IMARIS software on 3D reconstructed acini. Assuming that MCF10A cultured in 2D or in 3D react similarly to doxycycline, we determined the average number of GFP-NLP^+^ CRBs per GFP-NLP^+^ cell (= 2.09 ±0.1208 s.e.m.) by analyzing 2D cultured MCF10A (n=182 cells) treated with doxycycline for the same duration as used for 3D experiments. For each acinus, the number of GFP-NLP^+^ cells was then estimated by dividing the number of CRBs within the acinus by the average number of CRBs per NLP-GFP^+^ cell as determined above (= 2.09). Finally, the percentage of GFP-NLP^+^ cells within the acinus was then calculated by dividing the estimated number of GFP-NLP^+^ cells by the total number of cells within the acinus determined by DAPI staining.

### Western blots

Cells were harvested and proteins extracted in Extract buffer (50 mM Tris–HCl, pH 7.4, 0.5% IGEPAL (Sigma-Aldrich, St. Louis, MO, USA), 150 mM NaCl, 1 mM DTT, 5% glycerol, 50 mM NaF, 1 mM PMSF, 25 mM β-glycerophosphate, 1 mM vanadate, complete mini protease inhibitor cocktail (Roche Diagnostics, Basel, Switzerland). Proteins were then separated on Biorad mini-protean 4-15% gels (456-1086, Biorad Hercules, CA, USA) and transferred onto nitrocellulose membranes. Primary antibodies used were anti-E-Cadherin (1:5,000, 610182, BD, Franklin Lakes, NJ, USA) and anti-histone H3 (1:1,000, ab1791, abcam Cambridge, U.K) and secondary antibodies were HRP-conjugated anti-mouse immunoglobulin (170-6516, 1:2,000, Bio-Rad, Hercules, CA, USA) or anti rabbit immunoglobulin (170-6515, 1:2,000, Bio-Rad, Hercules, CA, USA).

### Mechano–optical microscopy

All AFM experiments were carried out under close to physiological conditions using a customized mechano-optical microscope (MOM) comprised of an AFM (JPK Instruments AG, Germany and SPECS Zurich GmbH, Switzerland) and epifluorescence/spinning disk confocal setup (Visitron Systems GmbH, Germany). Bright field and fluorescence images were taken with 10 and 40 X air objectives (Leica, Germany) and recorded using an ORCA Flash 4 sCMOS (Hamamatsu Photonics K.K., Japan). For AFM experiments, triangular DNP-S10 D (Bruker AFM Probes, USA) with a nominal spring constant of 0.06 N.m^−1^, an average tip length of 5 µm and a nominal tip radius of 15 nm, as well as rectangular HQ-CSC38/CR-AU B (MikroMasch, Nanoworld AG, Switzerland) cantilevers with a nominal spring constant of 0.03 N.m^−1^, and average tip length of 15 µm and a nominal tip radius of 20 nm were used. The experimental value for the spring constant of each cantilever was determined by thermal tune method (Sader et al., 1995). For cell stiffness measurements, the AFM was operated in a “Force-Volume Mode”. Briefly, arrays of force– displacement curves were recorded in a regular grid over a selected sample surface ranging from 20 × 20 μm to 80 × 80 μm and 32 × 32 to 64 × 64 force curves. Force–displacement curves were sampled with 4 kHz. All indentations were performed with the maximum load set to 1.8 nN at an indentation velocity of 16 μm.s^−1^. Individual AFM stiffness maps were completed within 20 to 45 minutes.

For measurements of different MDCK cell mutants (CEP68, PLK4, NLP; Dox-inducible over-expression) as well as for drug testing experiments, DNP-S10 D cantilevers were used. For 2D experiments, cells were plated on Ibidi μ-Dish 35mm, low with glass bottom or TPP 9.2cm^2^ dishes at 75,000 or 150,000 cells per cm^2^ and grown for 48 or 72 hours, respectively, to reach confluency. Doxycyclin was added at 2.5 μg.ml^−1^ for 24 hours to induce overexpression of NLP, CEP68 and PLK4. Before AFM measurements, confluent cells were rinsed with PBS and fresh doxycycline-containing growth medium was added to the cells. Drugs were added in the medium 30 minutes before AFM measurements; taxol (Merck Millipore Darmstadt Germany) was added at a final concentration of 50 nM, nocodazole (Sigma Aldrich MO, USA) at 200 ng.ml^−1^ and cytochalasin D at 10 μM.

For measurements of cells in 3D and comparison of mitotic versus interphase cells HQCSC38/Cr-Au B probes were used. Epifluorescence microscopy was used to identify GFP-CEP68, GFP-PLK4 and GFP-NLP over-expressing cells in 2D and GFP-NLP^+^ cells in 3D cultures. Furthermore, by using epifluorescence, mitotic cells were discriminated from interphase by visualization of mCardinal-H1 signal. Nanomechanical measurements of living 3D MDCK cysts were performed in culture dishes (TPP, Switzerland) previously coated with poly-L-lysine (P4707, Sigma Aldrich, USA) for 30 minutes and then rinsed with PBS before placing the cysts. For these experiments, individual cysts were first isolated from large pieces of embedding Matrigel with a pipette and placed in the culture dish. Cellular morphology and stiffness was used as an indicator that isolated cysts were free of Matrigel. All measurements were performed in medium containing 5% CO2 that was regularly replenished to maintain physiological conditions and compensate for evaporation.

Brightfield and fluorescence images were overlaid to identify WT and NLP over-expressing cells. The optical images were analyzed using Fiji/ImageJ (http://imagej.nih.gov). AFM data were further analyzed using custom-made “OfflineReader”, OriginPro 2016 (OriginLab Corporation, USA) as described previously (Plodinec et al., 2012; Plodinec et al., 2011) and Gwyddion (http://gwyddion.net/) software. AFM stiffness analysis over a wide range of conditions (2D, 3D, confluent, sub-confluent, impact of cytoskeleton drugs) was performed by normalizing the stiffness values of transgene-expressing cells to WT cells (non-overexpressing GFP-NLP), similarly to previously described analyses (Kerr et al., 2015). For individual maps in Fig 5A; the stiffness values of NLP over-expressing cells were first normalized against cells that did not overexpress centrosomal protein within one force map. For all other graphs, the stiffness values were directly normalized to the values obtained for WT cells in interphase. In all graphs, box plots show the mean (square), median (line), the standard error of mean (box) and the standard deviation (whiskers). Box plots show the mean (square), median (line), the standard error of mean (box) and the standard deviation (whiskers). For analyses, the statistical significance was tested using two-sample student’s t-test for normally distributed samples or otherwise using a Mann-Whitney test (Fig. 5B, 5D, Supp. Fig. 9C).

## Acknowledgments

We thank Inke Naethke (University of Dundee, UK), Channing Der (Pharmacology - UNC School of Medicine, USA), Daniel Gerlich (IMBA, Austria), Michael Davidson, Tilman Brummer (University of Freiburg), Stefan Zimmermann and Ralph Wäsch (University Medical Center Freiburg) for reagents. We also thank Elena Nigg for expert technical assistance, Kai Schleicher, Alexia Loynton-Ferrand, Wolf Heusermann and Frederik Grüll (imaging core facility) for assistance with imaging, and Janine Bögli (FACS core facility) for assistance with flow cytometry. Work in E.A.N.’s laboratory was supported by the Swiss National Science Foundation (310030B_149641) and the University of Basel. Work in the laboratory of R.Y.H.L was supported by the Canton of Aargau (Argovia Professorship awarded to R.Y.H.L), the Biozentrum, the Swiss Nanoscience Institute, and the Swiss National Science Foundation Nanotera Project awarded to the PATLiSci II Consortium.

## Author contributions

O.G. and D.S. contributed equally to this work. O.G. designed and analyzed most budding experiments, generated stable cell lines, performed and analyzed immunofluorescence and time lapse experiments, designed and performed E-Cadherin JSI experiments and determined and interpreted the estimated number of GFP-NLP^+^ cells per sphere. D.S. initiated the project and performed preliminary work, generated stable cell lines, designed and analyzed budding experiments, performed and analyzed invadopodia formation assays. P.O. designed, performed and analyzed AFM experiments. M.P. and R.Y.H.L. contributed to design and analysis of AFM experiments E.A.N supervised the project, designed experiments, analyzed the data and wrote the paper. All authors contributed to editing the manuscript.

## Conflict of Interest

The University of Basel has filed patents related to the AFM technology based on the inventions of M.P., P.O. and R.Y.H.L. Correspondence should be addressed to E.A.N.

